# Reduced representation sequencing accurately quantifies relative abundance and reveals population-level variation in *Pseudo-nitzschia* spp.

**DOI:** 10.1101/2022.05.06.490957

**Authors:** Carly D. Kenkel, Jayme Smith, Katherine A. Hubbard, Christina Chadwick, Nico Lorenzen, Avery O. Tatters, David A. Caron

**Affiliations:** Department of Biological Sciences, University of Southern California, 3616 Trousdale Parkway, Los Angeles, CA 90089, USA; Southern California Coastal Water Research Project, 3535 Harbor Boulevard, Suite 110, Costa Mesa, CA, 92626, USA; Florida Fish and Wildlife Conservation Commission-Fish and Wildlife Research Institute (FWC-FWRI), 100 8th Ave. SE, St. Petersburg, FL 33701, USA; Biology Department, Woods Hole Oceanographic Institution, 266 Woods Hole Road, Woods Hole, MA 02543, USA; U.S. Environmental Protection Agency, Gulf Ecosystem Measurement and Modeling Division, 1 Sabine Island Drive, Gulf Breeze, FL, 32561, USA

**Keywords:** harmful algal bloom, community composition, population genomics, next-generation sequencing, 2b-RAD

## Abstract

Certain species within the genus *Pseudo-nitzschia* are able to produce the neurotoxin domoic acid (DA), which can cause illness in humans, mass-mortality of marine animals, and closure of commercial and recreational shellfisheries during toxic events. Understanding and forecasting blooms of these harmful species is a primary management goal. However, accurately predicting the onset and severity of bloom events remains difficult, in part because the underlying drivers of bloom formation have not been fully resolved. Furthermore, *Pseudo-nitzschia* species often co-occur, and recent work suggests that the genetic composition of a *Pseudo-nitzschia* bloom may be a better predictor of toxicity than prevailing environmental conditions. We developed a novel next-generation sequencing assay using restriction site-associated DNA (2b-RAD) genotyping and applied it to mock *Pseudo-nitzschia* communities generated by mixing cultures of different species in known abundances. On average, 94% of the variance in observed species abundance was explained by the expected abundance. In addition, the false positive rate was low (0.45% on average) and unrelated to read depth, and false negatives were never observed. Application of this method to environmental DNA samples collected during natural *Pseudo-nitzschia* spp. bloom events in Southern California revealed that increases in DA were associated with increases in the relative abundance of *P. australis*. Although the absolute correlation across time-points was weak, an independent species fingerprinting assay (Automated Ribosomal Intergenic Spacer Analysis) supported this and identified other potentially toxic species. Finally, we assessed population-level genomic variation by mining SNPs from the environmental 2bRAD dataset. Consistent shifts in allele frequencies in *P. pungens* and *P. subpacifica* were detected between high and low DA years, suggesting that different intraspecific variants may be associated with prevailing environmental conditions or the presence of DA. Taken together, this method presents a potentially cost-effective and high-throughput approach for studies aiming to evaluate both population and species dynamics in mixed samples.

**Highlights:**

- 2bRAD method facilitates species- and population-level analysis of the same sample
- Method accurately quantifies species relative abundance with low false positives
- Consistent shifts in allele frequencies were detected between high and low DA years
- Certain *Pseudo-nitzschia* spp. populations may be more associated with DA presence

## 1. Introduction

Multiple species of the diatom genus *Pseudo-nitzschia* produce the water-soluble neurotoxin, domoic acid (DA). The toxin bioaccumulates via food web transfer, leading to illness and mass mortality of marine animals (Bejarano et al., 2008; Kvitek et al., 2008; Moriarty et al., 2021), significant economic costs to commercial and recreational shellfisheries (Moore et al., 2020; Wessells et al., 1995), and amnesic shellfish poisoning in humans who consume contaminated seafood (Trainer et al., 2012). Blooms of toxigenic *Pseudo-nitzschia* cause significant negative impacts in many regions of the world, including along the west coast of North America, where fishery closures and animal mortality occur frequently (Scholin et al., 2000; Smith et al., 2018a; Trainer et al., 2012). In California, blooms have been increasing in frequency and severity in recent years (Schnetzer et al., 2013; Trainer et al., 2010). In 2007, DA concentrations in mussels collected in California reached 610 µg g^-1^, more than 30 times higher than the U.S. Food and Drug Administration (FDA) tissue safety level of 20 µg g^-1^ (Trainer et al., 2012). In 2014, multiple mussel samples from Marin County contained DA concentrations in excess of 1000 µg g^-1^ (Langlois et al., 2014). *Pseudo-nitzschia* spp. bloom ‘hot spots’ in California are spread along the coast, from Monterey Bay in the north to San Luis Obispo and Point Conception, and the San Pedro/Long Beach Harbor area (Schnetzer et al., 2013; Smith et al., 2018a; Trainer et al., 2012), but it is not clear what attributes unite these geographically distant sites to promote HAB development.

Environmental factors implicated in bloom formation include nutrient enrichment from terrestrial sources, coastal upwelling, and mesoscale eddies (Anderson et al., 2008; Schnetzer et al., 2013). The grand majority of DA impacts on coastal ecosystems have been described in eastern boundary upwelling regions, including along the U.S. west coast where the California Current prevails (Trainer et al., 2010). Prior work has suggested that in California, blooms are associated with weak upwelling, lower salinity, temperature gradients and low macronutrient levels (Kudela et al., 2002). Culture and field studies have shown that phosphorus and silicate limitation can increase DA production (Fehling et al., 2004). The form of nitrogen can also play a role in DA production (Cochlan et al., 2008; Howard et al., 2007; Kudela et al., 2008), as can trace metal availability, including iron limitation and copper toxicity (Wells et al., 2005). Increased pCO_2_ has also been shown to increase DA production, and increased pCO_2_ in combination with silicate limitation has a synergistic effect on DA production (Tatters et al., 2012). In addition to the abiotic environment, biotic interactions, such as bacterial associations have also been implicated in toxin production (Bates et al., 1995). However, in field studies, consistent relationships between DA levels and specific environmental conditions have been less clear, with one study finding an association with colder, more saline water (Schnetzer et al., 2013); another, with dissolved silicic acid concentration (Smith et al., 2018b). While bloom risk mapping models have been developed for the California coast (C-HARM, (Anderson et al., 2016)), it remains challenging to predict the onset and severity of *Pseudo-nitzschia* bloom and toxic events, in part because the underlying drivers have not been fully resolved (Lelong et al., 2012; Smith et al., 2018b) and also because blooms are often a mixture of co-occurring species not all of which are toxigenic, or actively producing toxins for those that are.

More than 52 species of *Pseudo-nitzschia* have been described, but not all are toxigenic (Bates et al., 2018). Among the 26 species currently known to produce DA, toxin production was reported to be non-constitutive (Bates et al., 2018; Lelong et al., 2012; Trainer et al., 2012). Thus, variation in toxin concentrations during *Pseudo-nitzschia* blooms may stem from the presence or absence of exacerbating environmental factors and the ability to accurately measure such factors. This difference may also be due to an inability to accurately identify species- and population-level variation in *Pseudo-nitzschia* bloom composition. Recent work in several systems has indicated that the presence of certain species may be an important predictor of bloom toxicity, in addition to prevailing physiochemical factors such as temperature, salinity and nutrients (Clark et al., 2019; Smith et al., 2018b). In the San Pedro/Long Beach Harbor bloom hot spot, elevated DA concentration has been associated with community-level dominance of *Pseudo-nitzschia australis* and/or *P. seriata (Schnetzer et al*., *2013; Smith et al*., *2018b). P. multiseries* has also been reported as a major toxin producer in California (Trainer et al., 2000), but further north in Washington, *P*. cf. *pseudodelicatissima* and *P. cuspidata* join *P. australis* and *P. multiseries* as dominant toxin producers (Trainer et al., 2009).

Taxonomic identification of protists, such as *Pseudo-nitzschia*, is largely based on morphology, but salient features may not be sufficient to distinguish species with similar morphologies (Adl et al., 2007). Ultrastructural features of diatom cell walls visualized using scanning electron microscopy have long been the gold standard for species delineation (Round et al., 1990). However, the application of DNA sequencing revealed significant cryptic diversity – morphologically similar algae can have different genetic profiles (Alverson, 2008). Various genetic assays have been developed to genotype *Pseudo-nitzschia* spp., including direct sequencing of ribosomal gene regions (Casteleyn et al., 2010; Lim et al., 2014; Orsini et al., 2004), and DNA fingerprinting (Bornet et al., 2005) including microsatellites (Evans et al., 2004; Evans and Hayes, 2004). Automated Ribosomal Intergenic Spacer Analysis (ARISA) was more recently developed to distinguish *Pseudo-nitzschia* species within a heterogeneous bloom without the need for culturing (Hubbard et al., 2008), but this method still relies on a single locus, which limits the resolution of population genetic analyses because estimates of relatedness based on single loci are noisy (Lynch and Milligan, 1994). Furthermore, in spite of its broader field applicability, ARISA is still unable to fully resolve differences among certain *Pseudo-nitzschia* species in California, including *P. australis*, putatively one of the most toxic species, and another potentially toxic species, *P. seriata* (Hubbard et al., 2008).

Metagenomic analyses have the potential to provide increased taxonomic resolution, but as *Pseudo-nitzschia* spp. genomes can be large (e.g. *P. multiseries*, 218 Mbp, (Osuna-Cruz et al., 2020)) it will be challenging to cost-effectively obtain sufficient coverage for high-resolution spatial and temporal datasets. Reduced representation sequencing using Restriction site-Associated DNA (RAD) was originally developed for population genetic analyses of multicellular organisms (Hohenlohe et al., 2010). However, this method has the potential to cost-effectively facilitate understanding of both community and population-level variation in single-cell eukaryotes. RAD can identify and score thousands of genetic markers from individual or pooled samples (Davey and Blaxter, 2010; Schlötterer et al., 2014). It is analogous to other molecular genotyping methodologies, such as restriction fragment length polymorphisms (RFLPs) and amplified fragment length polymorphisms (AFLPs), which have been used to study phytoplankton population genetics since the late 1970s (Bruin et al., 2003), in that it reduces the complexity of the genome by subsampling only restriction enzyme recognition sites (Davey and Blaxter, 2010). However, RAD greatly surpasses these and other methods, including microsatellites, in its ability to identify, verify and score markers simultaneously and without an extensive developmental process – genotyping can be accomplished ‘*de novo*’, even on species with no prior sequence data (Davey and Blaxter, 2010; Matz, 2018). 2b-RAD is a type of RAD sequencing which uses type IIB restriction enzymes (Wang et al., 2012). These enzymes cut upstream and downstream of their recognition sites, generating uniform 36-bp DNA fragments, or tags, ideally suited for next-generation short read sequencing technologies. Analysis of within-species (∼population) genomic variation is based on identifying single nucleotide polymorphisms (SNPs) that occur within these 36-bp fragments and calculating allele frequencies based on read counts of tags containing alternate alleles. Here we develop a novel next-generation sequencing based assay using 2b-RAD and apply it to mock *Pseudo-nitzschia* communities generated by mixing cultures of different species in known abundances and field samples collected during natural *Pseudo-nitzschia* spp. bloom events.

## 2. Methods

### 2.1 Reference Cultures

*Pseudo-nitzschia* cultures were initiated from individual chains collected from multiple locations in the North Pacific. Seven strains representing four species, *P. australis* (isolated at Gaviota Beach, Santa Barbara County, California, 34.47° N, 120.23° W in April 2016), *P*. sp. (2 cultures, isolated from near Station Aloha 22.75°N, 158°W in May 2016), *P. pungens* (3 cultures, isolated from Newport Beach, Orange County, California, 33.61° N, 117.93° W in March 2017), and *P. subpacifica* (also isolated from Newport Beach, Orange County, California in March 2017), were grown in a modified F medium with 50% nutrient concentration (35.3 μM NO^3-^, 4.24 μM SiO_4_^2-^, 1 μM PO_4_^3-^) in autoclaved 0.2 µm filtered seawater. Incubators maintained cultures at 15°C, except for the *P*. sp. cultures which were maintained at 22°C. Cells were maintained under a 12h:12h light:dark cycle using cool white fluorescent illumination (∼110 μmol photons m^-2^ s^-1^). Cultures were visually inspected every 7-10 days with a dissecting scope for morphological signatures of health including proper cell shape and chain formation and transferred to new media on this same schedule.

*Pseudo-nitzschia* species were initially distinguished by overall size, shape of valve ends, and degree of valve overlap within chains by examination up to 400x magnification using light microscopy. *P. australis* (Fig. S1A), *P. pungens* (Fig. S1B) and *P. subpacifica* isolates were identified using frustule features including poroids, interstriae and fibulae observed via scanning electron microscopy (SEM). Respective samples were processed according to Hasle and Syvertsen (Hasle and Syvertsen, 1997). The cleaned frustules were observed with a Philips XL 30 S, FEG SEM. The small-celled *P*. sp. isolates were evaluated using light microscopy.

### 2.2 Generating Mock Communities

Mock communities were generated with cultured isolates identified to species level. Replicate community mixes with individual species representatives ranging in relative abundance from 10 (0.01%) to 99,990 cells (99.99%) in 100,000 total cells (Table 1) were generated to approximate the minimum bloom threshold of 80,000 cells/L previously described at Newport Pier, CA (Seubert et al., 2013). Triplicate cell counts were completed using a Sedgwick Rafter for each reference species after staining a 4-mL aliquot with 50 µL/mL Lugol’s Solution. A minimum of 400 cells were counted in each count and the relative standard deviation of the triplicate counts ranged from 2-10% (Table S1). Culture volumes were pooled within 4 hours of cell enumeration and immediately deposited onto 25mm GF/F filters (nominal pore size 0.7 microns) which were stored at -20°C until DNA extractions were completed. The level of detection for cellular enumeration was 3,000 cells/L for each individual count and so mix fractions represented by less than 3,000 cells (0.1 and 0.01%) were made through serial dilution prior to mixing. There were 6 mix types in addition to pure culture samples as detailed in Table 1.

**Table 1.** Replication of mock culture sample mixing scheme.

| Targeted cell<br>abundance ratio | Percent<br>abundance ratio | N replicate<br>mixes per ratio |
| --- | --- | --- |
| 100,000 | 100% | 3-5 per species |
| 99,990 : 10 | 99.99% : 0.01% | 8 |
| 99,900 : 100 | 99.9% : 0.1% | 4 |
| 75,000 : 20,000 : 5,000 | 75% : 20% : 5% | 3 |
| 66,000 : 33,000 | 66% : 33% | 1 |
| 60,000 : 30,000 : 10,000 | 60% : 30% : 10% | 4 |
| 40,000 : 30,000 : 30,000 | 40% : 30% : 30% | 4 |
| 33,000 : 33,000 : 33,000 | 33% : 33% : 33% | 3 |
| 50,000 : 50,000 | 50% : 50% | 3 |

### 2.3 DNA extraction from mock community mixes

Pure culture and mock community mix samples were extracted using a phenol-chloroform protocol modeled after (Countway et al., 2005). Briefly, 2 mL of lysis buffer comprised of 40 mM EDTA (ph 8), 100 mM Tris (ph 8), 100mM NaCl, and 1% SDS weight/volume was added to frozen filters in 15 mL Falcon tubes. Tubes were thawed at 70°C in a water bath after which 200 µL of 0.5 mm zirconia/silica beads (BioSpec Products) were added. The mixture was vortexed for 1 minute then heated at 70°C for 5 minutes for four repetitions to lyse the cells and break down the filter. The lysate and filter mixture was transferred to a 10 mL syringe and lysate was dispensed into a new 15 mL Falcon tube. 2.5 M NaCL and 10% CTAB were added to yield a 0.7 M NaCl 1% CTAB solution and the full mixture was incubated for 10 minutes at 70°C.

An equal volume of 25:24:1 phenol:chloroform:isoamyl alcohol (Sigma) was added and the mixture was vortexed and centrifuged at 4750 RPM for 10 minutes. After the layers settled the bottom layer was removed via pipetting and discarded. An equal volume of phenol:chloroform:isoamyl alcohol was again added and the mix was vortexed and centrifuged at 4750 RPM for 10 minutes. The supernatant was removed and placed in a new Falcon tube.

The supernatant was extracted by performing two repetitions of adding an equal volume addition of 24:1 chloroform: isoamyl alcohol (Sigma), vortexing, centrifuging at 4750 RPM for 10 minutes and discarding the bottom layer up to the interface. The supernatant was distributed into 1.5 mL Eppendorf tubes in 400 µL aliquots. Each aliquot was precipitated in 800 µL of cold 95% ethanol and 0.1X volume of 10.5 M ammonium acetate (40 µL), vortexed and incubated overnight at -20°C.

Following incubation samples were centrifuged at 14000 RPM for 30 minutes at 4°C. Liquid was decanted and the pellet was rinsed in 70% cold ethanol and centrifuged at 14000 RPM for 15 minutes at 4°C after which the liquid was decanted and the pellet was allowed to air dry. The pellet of each aliquot was suspended in 30 µL MilliQ H_2_O and aliquots from the same sample were combined for DNA quantitation.

Extractions were further purified using a Genomic DNA Clean & Concentrator-10 kit (Zymo Research) according to the manufacturer’s instructions. Briefly, the eluted DNA was combined with buffer and placed into a spin column. Samples were washed and centrifuged before being eluted in 10 µL elution buffer. Samples were vacuum-concentrated using a Speed Vac (Thermo Fisher, USA) on low speed for 1 min to yield a sufficient volume for the 2bRAD protocol (∼8 µL).

### 2.4 Natural Time-series Samples

Weekly water samples were obtained as part of long-term SCCOOS & CeNCOOS Harmful Algal Bloom Monitoring Alert Program (https://sccoos.org/harmful-algal-bloom/) at Newport Beach Pier, CA (Kudela et al., 2015; Seubert et al., 2013; Smith et al., 2018a). Samples have been routinely collected since 2008 for the quantification of domoic acid and enumeration of harmful phytoplankton taxa, as well for basic physicochemical conditions. Particulate domoic acid (pDA) samples were collected via the filtration of 200 mL of sea water onto a GF/F filter and were frozen at -20°C until extraction. Samples were quantified via Enzyme-Linked ImmunoSorbant Assay (ELISA: Mercury Science, Durham, NC) with a detection limit of 0.02 ng/mL. Samples below the detection limit were assumed to be zero for all calculations and statistical analyses. Seawater samples were collected and preserved with 3% formaldehyde (final concentration) for the enumeration of harmful phytoplankton taxa including *Pseudo-nitzschia*. Cells were enumerated using a Leica DM IRBE inverted light microscope (Leica Microsystems, Buffalo Grove, IL) at 400x after settling 25 mL of the sample in Utermöhl chambers for approximately 24 hours (Utermöhl, 1958). *Pseudo-nitzschia* cells were categorized into seriata size class (> 3 µm) and delicatissima (< 3 µm) size classes. Light microscopy based size sorting was conducted to differentiate between larger species more often associated with detectable pDA in the region and smaller cells that are more rarely associated with high toxin concentrations, as described in Seubert et al. (2013).

DNA was extracted from archived HABMAP sampling filters for 2bRAD sequencing. Samples were selected from the spring bloom in four different years in which *Pseudo-nitzschia* spp. cells were quantified, two in which in pDA was observed (2017 and 2019) and two in which no DA was detected (2015 and 2018). Briefly, 500 mL of water were filtered onto a 25mm GF/F filter and subsequently frozen at -20°C. DNA was extracted from filters using a Qiagen Plant kit (Qiagen, USA).

### 2.5 Library Preparation and Sequencing

DNA samples from the mock community mixes were prepared for 2bRAD sequencing using the BcgI restriction endonuclease targeting 100% of restriction sites (Wang et al., 2012). The same protocol was used for the natural time-series samples, save that a 1/64 site reduction scheme was used. The full library preparation protocol is available at https://github.com/z0on/2bRAD_denovo. Culture reference and mock community mix libraries were run on two lanes of Illumina HiSeq 2500, SR 50 format. Natural bloom libraries were run on four lanes of Illumina Nextseq 550, SR 75 format. All sequencing runs were carried out by the University of Southern California Genome core.

### 2.6 Generation of RAD reference libraries

All reads originating from pure culture samples were used to create independent RAD tag reference libraries for subsequent read mapping. 2bRAD tag references were also generated for the *Pseudo-nitzschia multiseries* reference genome (CLN-47, JGI) and a whole genome shotgun sequence set for *Pseudo-nitzschia multistriata* (CAACVS01, GCA_900660405, ENA). Full scaffolds, mitochondria, and plastid assemblies were concatenated and a custom perl script was used to extract tags exhibiting the Bcg I restriction site motif. Full bioinformatic protocols and scripts can be found at https://github.com/ckenkel/Pseudo-nitzschia2bRAD. Briefly, for sequenced cultures, a custom perl script was used to trim sequencing adapters from raw reads, retaining only the 36-bp insert. These trimmed files were quality filtered using the fastx_toolkit (Assaf and Hannon, 2010) requiring a minimum phred quality of 20 over 100% of the read. The bbduk command of the BBMap package (Bushnell, 2014) was then used to filter reads matching to Illumina sequencing adapters. Reads passing these filtering criteria were clustered at 100% identity using a custom perl script to remove perfect duplicates, and only reads exhibiting a 100% match to the restriction site were retained (Table S1).

As cultures were not axenic, reads were then filtered for contaminants based on a BLAST search against the NCBI nt database ((Sayers et al., 2019), downloaded 17 Jun 2020). Up to five alignments were reported for each read for matches below the e-value threshold of 10^−5^. A custom perl script was then used to summarize best hits using the NCBI taxonomy (Schoch et al., 2020). Contaminant reads were flagged as those exhibiting significant similarity to any non-Bacillariophyte sequence. Reads for each species were then clustered at 91% identity, which allows for up to 3 mismatches, or SNPs, per tag, using cd-hit (Fu et al., 2012; Li and Godzik, 2006). Species clusters containing any reads previously identified as matching to a contaminant were removed using a custom bash script. Species specific references and known contaminants were then concatenated into a ‘global’ reference, retaining a species-level identifier within each tag.

### 2.7 Analysis of mock community mixes and natural time-series samples

All bioinformatic and statistical scripts can be found at https://github.com/ckenkel/Pseudo-nitzschia2bRAD. A custom perl script was used to filter and trim raw reads. Reads were discarded if they did not contain the correct restriction site motif or adapter sequence, and those exhibiting the expected sequence construct were subsequently trimmed to remove adapter sequences. In addition, PCR duplicates were discarded by retaining only one representative read for sequences sharing the same 64-fold degenerate adapter sequence, the first 34 bases of the insert, and the secondary barcode. Adapter trimmed reads were further filtered for quality, retaining only reads exhibiting Q20 over 100% of bases and exhibiting no match to Illumina adaptors using the fastx_toolkit (Assaf and Hannon, 2010) and BBMap (Bushnell, 2014)(Table S2). Reads were mapped against the global species reference library using bowtie2 (Langmead and Salzberg, 2012), default parameters, including the --very-sensitive flag) and a custom bash script was used to count the number of reads exhibiting high quality matches (reads with MAPQ<23, discarding any ambiguous matches) to each species reference.

Samples with fewer than 1000 high quality mapped reads remaining were removed. The false positive rate (FPR) was calculated as the sum of reads matching species known to be in each sample relative to the total number of high quality mapped reads for that sample. Accuracy was calculated as the absolute value of the difference between observed and expected percent abundance values for each focal species within each sample.

Coverage of natural time-series sample reads mapping to *Pseudo-nitzschia* spp. in the global reference library was lower than the 80-100x minimum coverage recommended for pooled sequencing (Schlötterer et al., 2014). Therefore, to achieve sufficient read depth to assess allelic diversity within species over time, mapped reads in SAM files were split by species and concatenated by year. Tags with three or more SNPs were excluded from further analysis and remaining SNPs were thinned to one per tag. A Cochran-Mantel-Haenszel test as implemented in Popoolation2 (Kofler et al., 2011) was subsequently used to test for consistent changes in allelic diversity between high DA and low DA years. We required the minimum allele count to be 12, and the minimum and maximum coverage to be 50x and 200x respectively. P-values were calculated from pairwise comparisons of 2015 vs 2017 and 2018 vs 2019. Plotting and statistical analyses were carried out using the R language environment (v4.1.0, (R Core Team and Others, 2017).

### 2.8 Amplification and identification using ARISA

*Pseudo-nitzschia* species community composition in natural time series samples were also analyzed via ARISA. Genomic DNA concentrations were quantified on a plate reader (Biotek Instruments) using Picogreen (Invitrogen) and then standardized to 1ng µL^-1^ prior to amplification. Genus-specific oligonucleotides PnAll F (5′-TCTTCATTGTGAATCTGA-3′) and FAM-labeled PnAll R (5′-CTTTAGGTCATTTGGTT-3′) were used to amplify the ITS1 region (Hubbard et al., 2014, 2008). For PCR, 10 ng of genomic DNA was added to replicate (duplicate or triplicate) 20 μL reactions consisting of 2.5 mM deoxynucleoside triphosphates, 0.4 mM of each primer, 0.75 U of Apex Taq polymerase, 2 mM of MgCl_2_, and 1 x standard reaction buffer (Apex Bioresearch Products). Amplification was conducted using a 2–minute denaturation step at 94°C, followed by 32 cycles of 30 seconds at 95°C, 30 seconds at 50.6°C, and 60 seconds at 72°C, and ending with a 10 minute extension at 72°C. Resulting products were purified using MultiScreen PCRμ96 filter plates (Millipore). Replicate reactions were pooled, quantified using Picogreen, and diluted to 1 ng µL^-1^. For fragment analysis, 1 ng of PCR product was processed using an Applied Biosystems 3730 XL DNA Analyzer (University of Illinois DNA Core Sequencing Facility) with a LIZ600 size standard. Electropherograms were analyzed using DAx software (Van Mierlo Software Consultancy) to determine peak height and size (in base pairs or bp). Relative abundance of each amplicon was calculated by dividing the height of each individual peak by total peak height, and only those peaks that exceeded 3.0% of the total peak height were used in the final dataset following (Hubbard et al., 2014, 2008). Furthermore, only samples with a total peak height of >1000 relative fluorescent units (RFUs) were used. Relative peak height was previously determined to be correlated with the proportion of ITS1 copies added to the PCR, recognizing that larger-celled species generally have larger genomes and more ITS1 copies per cell (Hubbard et al., 2014) and was thus used to provide semi-quantitative data about changes in species contributions over time.

For each peak, 4 bp were added manually such that amplicon sizes for ARISA matched those based on *in sillico* sequence comparisons. A subset of samples was selected for confirmatory ITS1 sequencing where 1 μL of genomic DNA was amplified as described above (with an annealing temperature of 67°C) using 18SF-euk (5′-CTTATCATTTAGAGGAAGGTGAAGTCG-3′) and 5.8SR-euk (5′-CTGCGTTCTTCATCGTTGTGG-3′) oligonucleotides. The resulting PCR product was run on a 3% agarose gel. Using sterile micropipette tips, distinct bands were individually picked from the gel and then gel picks were dissolved in 10 μL of sterile molecular grade water for 5 minutes at 40°C prior to PCR amplification. The PnAll F/R primer set was utilized to amplify 1 μL of the melted gel/sterile water mixture and the resulting PCR product was run on a 3% agarose gel to confirm the presence of a single band and assess approximate product size; gel picks from these products were used in another round of PCR with the PnAll F/R primer set. A 1.5% agarose gel was used to confirm single band product sizes and remaining products were purified using ExoSap-It Express PCR Product Cleanup Reaction (Applied Biosystems). Purified products were then sequenced on a 3730 XL DNA Analyzer (Applied Biosystems) by Eurofins Genomics LLC. Sequences were analyzed using Sequencher software (Gene Codes Corporation) and identified using BLASTn to query the National Center for Biotechnology Information’s (NCBI) GenBank nucleotide database (Sayers et al., 2019) to verify sizes determined by ARISA. Linear models were used to compare the relative abundance of focal species derived using the different methods (ARISA vs 2bRAD) and a Fisher’s exact test was applied to compare presence/absence based detection using a contingency table comparing the overlap between positive and negative calls for each method.

## 3. RESULTS

### 3.1 Reference Library Construction

Sequencing yielded a total of 133,370-452,059 raw reads for each species (Table 2). Of these, between 6.5-24.4% were identified as non-Bacilliariophyte contaminants. The majority of contaminants exhibited best matches to Proteobacteria (Fig. S2). Some bacteria were previously identified as putative associates of diatoms and *Pseudo-nitzschia* spp. including *Sulfitobacter pseudonitzschiae*, which comprised 0.1-0.5% of the total contaminant sequences identified across species and *Marinobacter salarius*, which occurred at low abundance in most species (<5%) but dominated contaminant sequences in *P. australis* cultures, comprising more than 93% (76,814) of the tags identified as contaminants. Removal of any clusters with individual sequences exhibiting matches to contaminants yielded a final set of 71,889-113,898 read clusters by species, on par with the number of BcgI restriction sites identified in the *P. multiseries* reference genome (81,121, Table 2).

**Table 2.**
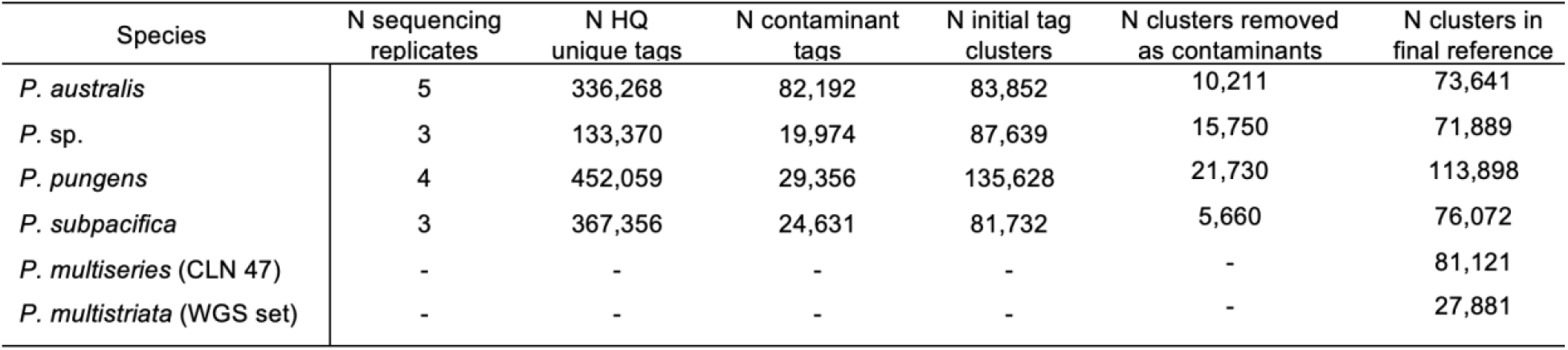
Sequencing effort, clustering, and contaminant removal to generate 2bRAD reference libraries per species. Tags: reads trimmed to retain only the BcgI restriction site motif; HQ: high quality, defined as reads exhibiting a minimum phred quality of 20 over 100% of the read and not containing any Illumina adapters.

### 3.2 Accuracy and precision of reduced representation sequencing based estimates of species abundance

The false positive rate (FPR), reflecting samples in which reads mapped to a reference species not included in that mock community mix, was low, 0.45% on average, and was unrelated to read depth (Fig. 1a). There was a significant difference in the FPR between mix types, with samples derived from multi-species pools exhibiting an 0.68% FPR on average (range: 0.045 - 4.1%) whereas the FPR of ‘pure’ culture samples was 0.04% on average (range: 0 - 0.1%, t(26.15)=3.48, p=0.0018, Fig. 1b). The highest false positive rates were observed in the most extreme mixes (ratios greater than 99:1, Fig. 1). False negatives were never observed. That is to say, species known to be included in the sample mix were always detected, even at very low relative abundances, e.g. 0.01% (Fig. 1).

**Figure 1.**
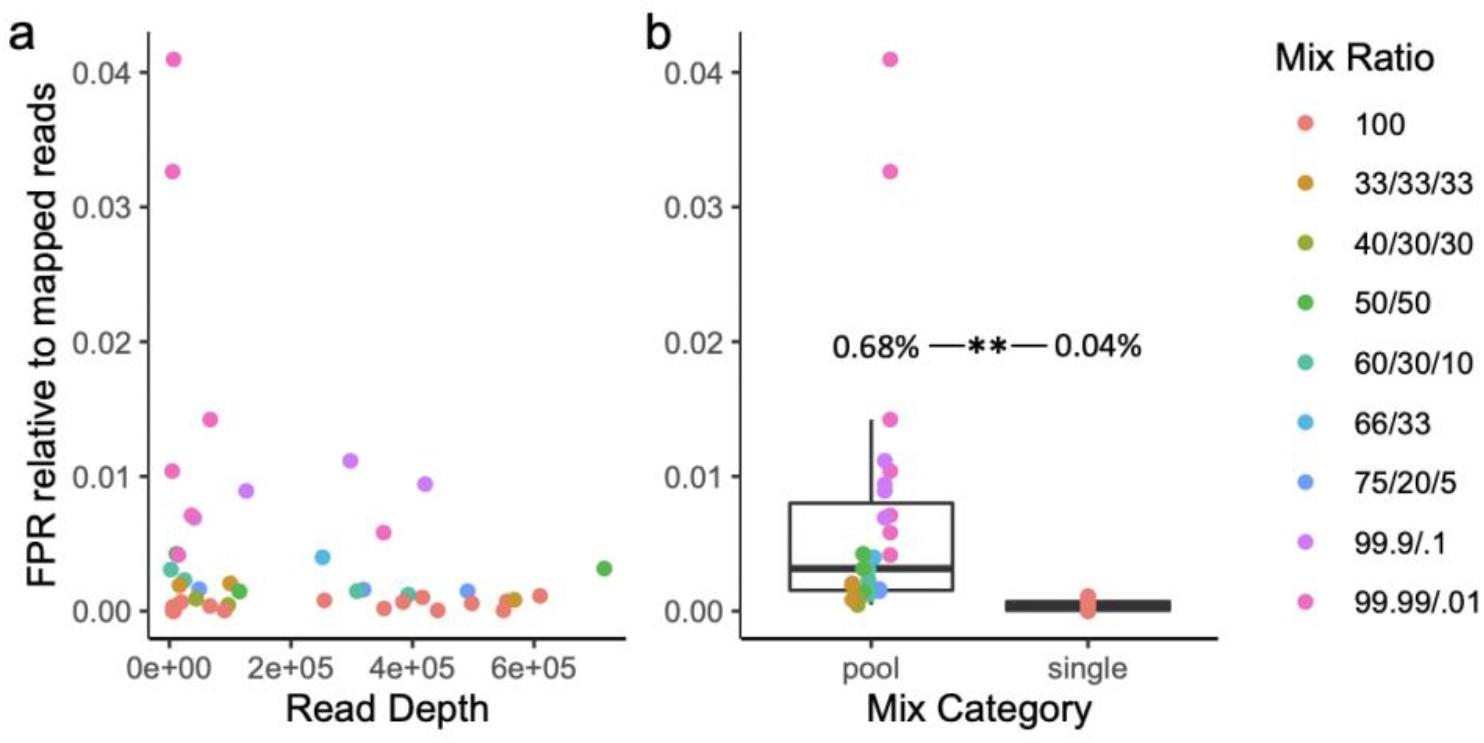
False positive rate (FPR) relative to high quality mapped reads as a function of read depth (a) and whether the sample consisted of a single species or multi-species pool (b). Individual samples are colored according to their mix type ratio (Table 1).

This pattern was also apparent when examining FPR by species, with significant increases in the FPR for reads mapping to *P*. sp. (t(26.1)=10.5, p=4e-11), *P. multiseries* (t(39.1)=2.5, p=0.008), *P. multistriata* (t(26)=2.89, p=0.004), and *P. pungens* (t(4.05)=4.45, p=0.005) references in multi-species mixes relative to individual species samples and similar trends for the remaining species (Fig. S3).

On average, 94% of the variance in observed species abundance was explained by the expected abundance based on the target mix ratios (R2=0.94, F(1,79)=1350, p<0.001, Fig. 2A). Expected abundances were also a strong predictor of observed abundances, and the intercept was not significantly different from zero (α=0.24, β=0.95, t(79)=36.7, p<0.001). However, fit did differ between species, with *P. australis* having a higher intercept on average than *P. pungens* and *P. subpacifica* (Fig. 2a). Accuracy was not a function of species (F(3,77)=2.45, p=0.07), although mixes with *P. subpacifica* tended to be more accurate (Fig. 2b). Accuracy did differ among mix types (F(8,72)=12.54, p=4.3e-11, Fig. 2c). More equal mixes, for example those with a ratio of 33:33:33 of three species, were less accurate than more extreme mixes, or those with a ratio >99:1 of two species (Tukey’s HSD < 0.001, Fig. 2c). Accuracy also differed among culture replicates (F(8,72)=4.38, p=0.0002, Fig. 3d). For example, more error was observed for the C5 culture replicate of *P. pungens* than for C3 (Tukey’s HSD=0.05, Fig. 2d).

**Figure 2.**
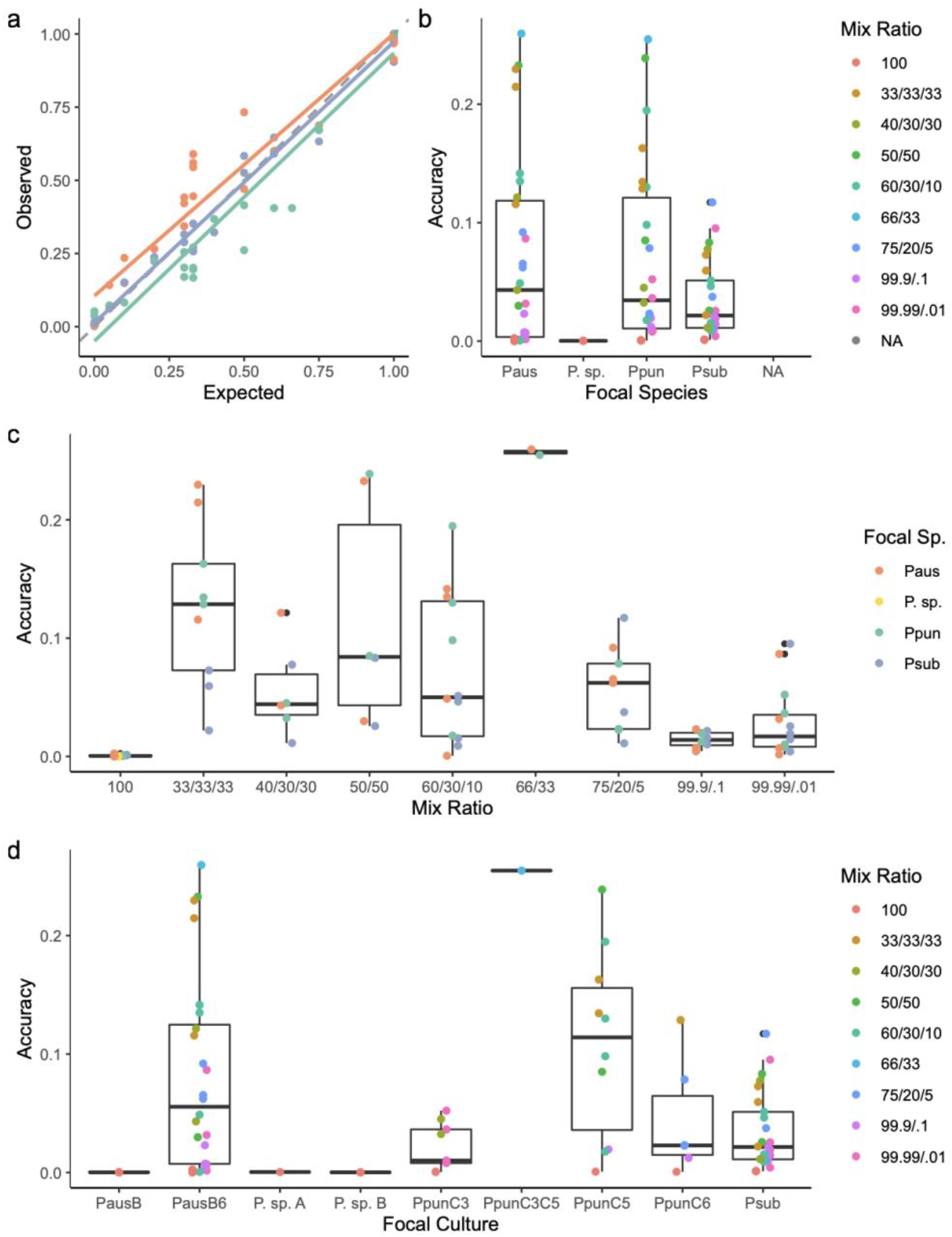
Accuracy of a reduced representation sequencing (2bRAD) method for quantifying species relative abundance. (a) Observed abundance, calculated as the proportion of reads mapping to the reference library, as a function of expected abundance based on the target proportion in the mock community mix varies by species. The dashed gray line is the 1:1 line. (b) Accuracy, or the deviation between observed and expected values, as a function of species and mix ratio. (c) Accuracy as a function of mix ratio and species. (d) Accuracy as a function of culture and mix ratio.

**Figure 3.**
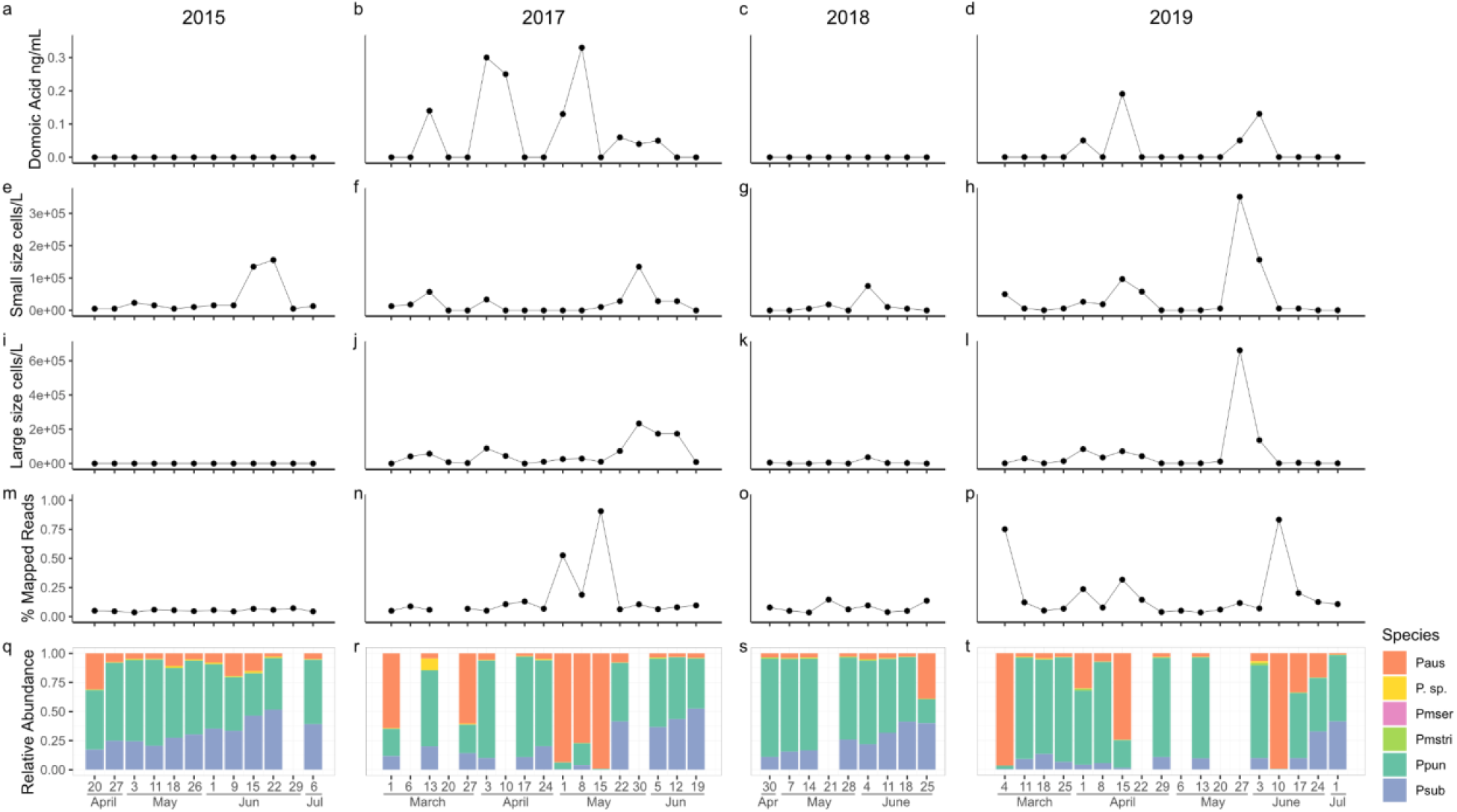
Dynamics of *Pseudo-nitzschia* spp. blooms sampled from Newport Beach Pier, CA. (a-d) The concentration of particulate domoic acid (ng/mL) in seawater. (e-h) The concentration of small size class (< 3 µm) cells per liter. (i-l) The concentration of large size class (> 3 µm) cells per liter. (m-p) The percent of high quality reads exhibiting high quality mapping to the *Pseudo-nitzschia* spp. reference library. (q-t) The relative abundance of *Pseudo-nitzschia* spp. calculated as the proportion of high quality reads exhibiting high quality mapping to each individual species relative to the total number of reads mapping to the *Pseudo-nitzschia* spp. reference library.

### 3.3 Abundance estimate of species in natural *Pseudo-nitzschia* spp. blooms using 2b-RAD

The dynamics of *Pseudo-nitzschia* spp. blooms differed across years. *Pseudo-nitzschia* spp. cells were observed every year, but pDA was only detected in 2017 and 2019 (Fig. 3a-c). Although samples were sequenced for 55 out of 56 total time-points, the overall percent of high quality reads mapping to the six species reference was generally very low (Fig. 3d). Consequently, we were unable to quantify species composition in some samples due to insufficient read depth (Fig. 1e, empty columns). Superficially, increases in pDA appeared to be associated with increases in the relative abundance of *P. australis*. However, the correlation across time-points was weak (R2=0.05, t(44)=1.8, p=0.08, Fig. 3,S4). The relative abundance of *P. australis* did explain a significant portion of the variation in the percent of reads mapping to the six species reference (R2=0.69, β=0.56, t(44)=10.06, p<0.001, Fig. S5).

### 3.4 Abundance of species in natural *Pseudo-nitzschia* spp. blooms using ARISA

Seventeen distinctly-sized amplicons were observed with ARISA (Fig. 4). Ten were associated with species previously reported along the US Pacific Coast based on reference sequences available in GenBank and prior studies using ARISA (Carlson et al., 2016; Hubbard et al., 2014, 2008; Smith et al., 2018b). Half were putatively identified based on prior observations as *P. australis/P. seriata* (which share the same fragment size, 150 base pairs [bp]), *P. inflatula* (156 bp), *P. delicatissima* (168 bp), *P. fraudulenta* (203 bp), and *P. fryxelliana* (207 bp). The other half were associated with species based on prior observations and direct sequencing of PCR products as part of the present study: *P. galaxiae* (140 bp), *P. pungens* (142 bp), *P. multiseries* (144 bp), *P. subpacifica* (196 bp) and *P. cuspidata* (233 bp); all were 100% similar to previously observed sequences. The sequence obtained for *P. pungens* corresponded to *P. pungens* var. *cingulata*. Seven additional amplicons of unknown identity were characterized by ARISA only (170 bp, 177 bp, 200 bp, 210 bp, 213 bp, 219 bp, and 223/224 bp). A few samples (n=3) with low total peak height were excluded from analysis (indicated by * in Fig. 4).

**Figure 4.**
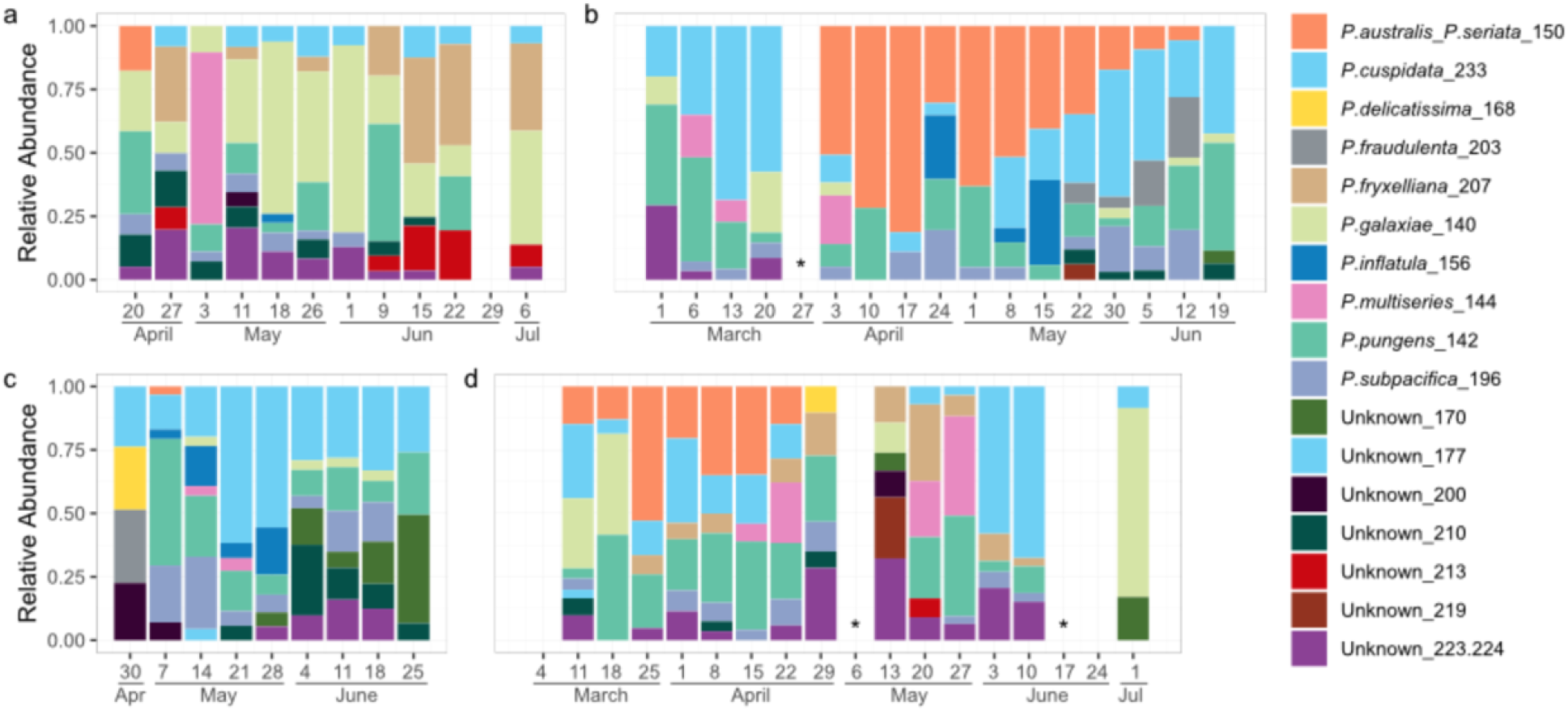
ARISA-based diversity profile of *Pseudo-nitzschia* spp. blooms sampled from Newport Beach Pier, CA in (a) 2015, (b) 2017, (c) 2018, and (d) 2019. * samples with low total peak height.

The *P. australis/P. seriata* fragment was observed in spring in all four years but the duration varied from year to year: 2015 (April, one week), 2017 (April-June, 11 weeks), 2018 (May, one week), and 2019 (March through April, seven weeks, Fig. 4). *Pseudo-nitzschia* cellular abundances and pDA concentrations generally tracked well with *P. australis/P. seriata*. For example, in 2015 and 2018, cellular abundances and pDA concentrations were minimal, and increases in the former only were noted after the single observations of *P. australis*/*P. seriata*. In contrast, in 2017 and 2019, cellular abundances and pDA concentrations were elevated and dynamic for the time frame when the *P. australis/P. seriata* amplicon was present, and even in a few samples where this amplicon was absent but other toxic species like *P. cuspidata* were present (Fig. 3,4). *P. cuspidata* was observed in nearly every sample, but appeared to be quite dynamic based on the relativized ARISA signal, and was detected before, during, and after *P. australis/P. seriata* in all years except 2015, the year that *P. cuspidata* generally appeared to comprise a smaller proportion of the *Pseudo-nitzschia* assemblage and when *P. galaxie* was more prevalent. *P. multiseries* is another species recognized as producing high pDA levels and was detected in all years, but was never the dominant taxon in the ARISA. *P. multiseries* was observed after *P. australis* in 2015, before and with *P. australis* in 2017, after *P. australis* in 2018, and with and after *P. australis* in 2019.

Although both methods identified an increase in *P. australis* in the high domoic acid years (2017 and 2019), correlations between relative abundance estimates for individual species obtained using the different methods were weak or nonexistent. A significant correlation was detected between the relative abundance of *P. australis* estimated via 2bRAD and the ARISA-based estimate of *P. australis/P. seriata*, but the overall variance explained was less than 10% (R2=0.096, t(40)=2.3, p=0.03, Fig. S6). No relationships were detected for any of the other species that could be compared between methods (*P*. sp., *P. pungens, P. multiseries, P. subpacifica*) nor were any relationships evident when converting relative abundance to simple presence/absence calls.

### 3.6 Population-level variation in *Pseudo-nitzschia* spp. over time

The low percentage of reads mapping to the *Pseudo-nitzschia* spp. reference in general (Fig. 3) resulted in insufficient coverage for SNP calling in individual samples. For example, even for *P. pungens*, the most abundant species across sampling time-points and years, 75% of samples had fewer than 7 tags with at least 80x coverage, the minimum recommended threshold for analysis of pooled samples (Schlötterer et al., 2014) (Fig. S7). Therefore, to assess the potential for the 2bRAD method to yield information on population-level variation, we pooled samples by year and compared changes in allele frequency between low domoic acid (DA, 2015 and 2018) and high domoic acid years (2017 and 2019, Fig. 3) following (Bendall et al., 2016).

The relative abundances of *P. delicatissima, P. multiseries*, and *P. multistriata* were too low to conduct population-level analyses with confidence (Fig. 3). No consistent changes in allele frequencies were detected in *P. australis* populations over time. We identified 17 SNPs in *P. pungens* and 4 SNPs in *P. subpacifica* that showed differences in allele frequencies across years (CMH<0.05). Of these, 10 in *P. pungens* and all of the *P. subpacifica* SNPs exhibited consistent changes between years with low and high pDA concentrations. In *P. subpacific*a, the minor allele frequency (MAF) at all loci consistently decreased in years where pDA was observed whereas in *P. pungens*, MAF decreased at 7 loci and increased at 3 loci (Fig. 5, Table S3).

**Figure 5.**
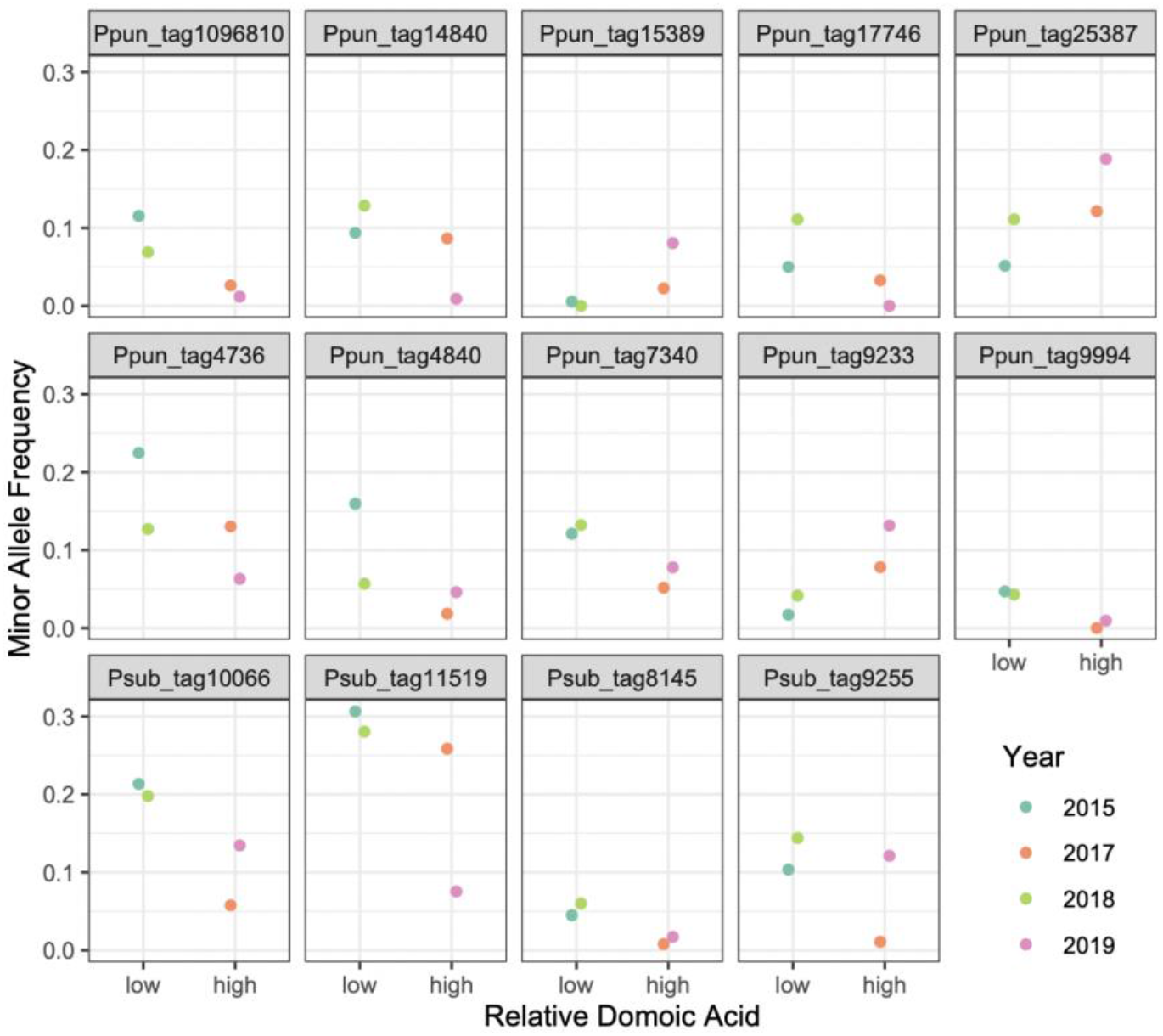
Minor allele frequency at select loci (CMH<0.05) by sampling year, stratified by the relative amount of domoic acid detected over the course of the sampling window. Ppun = *P. pungens*, Psub = *P. subpacifica*.

## 4. DISCUSSION

In culture, the toxicity of *Pseudo-nitzschia* is influenced by both abiotic (Trainer et al., 2012) and biotic (Sison-Mangus et al., 2014) factors. In the Southern California Bight, while large-scale drivers of domoic acid outbreaks have been identified, the factors contributing to interannual variability in toxin production remain unresolved (Smith et al., 2018a). Inter- and intraspecies level variation in *Pseudo-nitzschia* is thought to play a key role in DA production (Clark et al., 2019; Fernandes et al., 2014; Guannel et al., 2015; Hubbard et al., 2014; Smith et al., 2018b). Here we show that a reduced representation based sequencing approach can be used to accurately quantify the relative abundance of *Pseudo-nitzschia* spp. in a mixed community. In addition, we were able to assess population-level genomic variation within focal *Pseudo-nitzschia* spp. using the same dataset. This method presents a potentially cost-effective approach for large-scale studies aiming to evaluate population and community dynamics in mixed samples, although additional development work is needed.

### 4.1. The dynamics of Pseudo-nitzschia blooms at Newport Beach Pier, CA

Analysis of a field sample set over high and low domoic acid years suggest a role for *P. australis* in DA production. However, the direct linear relationship between the relative abundance of *P. australis* as estimated through 2bRAD and pDA concentrations is weak, although corroborated by ARISA, suggesting that the toxicity of *P. australis* is not a linear function of its abundance, or that other members of the *Pseudo-nitzschia* spp. community or variability in their abundances may be involved. The parallel ARISA dataset generated for the same samples supports this latter explanation: in addition to the high prevalence of *P. australis*/*P. seriata*, additional toxigenic or suspected toxigenic species such as *P. cuspidata, P. pungens, P. multiseries, P. fraudulenta, P. delicatissima, P. subpacifica and P. galaxiae* were also identified. An earlier ARISA based analysis of relative species abundance also identified *P. australis/P. seriata* in association with a strong DA event in 2013, but these species were absent in a similar bloom the following year (Smith et al., 2018b). ARISA cannot distinguish *P. australis* from *P. seriata* on the US west coast (Hubbard et al., 2008). Although *P. seriata* was not included in our reference library, the 2bRAD method is highly specific in terms of read recruitment and false positive rate, and given the strength of read recruitment to the *P. australis* reference when this species is dominant (Fig. S4), it is likely that *P. australis* rather than *P. seriata* is the dominant member of the *Pseudo-nitzschia* spp. assemblage for the years analyzed here, as false positives between sister species using the 2bRAD method, such as *P. pungens* and *P. multiseries*, are minimal (∼0.005%). If we can interpret the overall percent mapping as a reflection of the total composition (Fig. 3m-p), both for species within our *Pseudo-nitzschia* reference set and others, then *P. australis* may indeed dominate blooms in this region at certain sampling times and is often dominant when DA is detected. Indeed, both ARISA and 2bRAD detected dynamic assemblages over time and a coincident increase in *P. australis* as DA increased. However, an expansion of the 2bRAD reference library set and additional work will enable this supposition to be tested.

Interestingly, we also identified apparent cyclical shifts in allele frequency in both *P. pungens* and *P. subpacifica* populations wherein the minor allele frequency increased or decreased corresponding with high vs low DA years (Fig. 5). Prior temporal analyses of population level variation in SNPs from natural bacterial populations have identified unidirectional shifts in allele frequency over time (Bendall et al., 2016). Population genetic analysis of *Pseudo-nitzschia* spp. has been limited to microsatellite-based characterization (Casteleyn et al., 2010). Globally sampled populations of *P. pungens* were found to exhibit strong isolation by distance, however, time-series samples were pooled by region (Casteleyn et al., 2010), potentially masking relevant temporal shifts which have been repeatedly identified in other marine diatoms (Godhe and Härnström, 2010; Rynearson et al., 2006; Whittaker and Rynearson, 2017). Interestingly, morphologically and genetically distinct *P. pungens* varieties have been identified along the US West Coast (Carlson et al., 2016; Hubbard et al., 2014, 2008; Villac and Fryxell, 1998). Here we show that not only do allele frequencies show temporal shifts, in some instances these shifts correlate to some extent with the presence of DA (Fig. 5). It is unlikely that the SNPs identified here are causal, in the sense that they are responsible for DA production, but it is possible that certain *Pseudo-nitzschia* spp. populations are more associated with DA presence and/or production than others. The presence of competitors can alter both ecological and evolutionary outcomes (Agrawal et al., 2012). A competitive role for DA has been proposed, but the mechanism of action remains debated. DA has been hypothesized to play a role in iron uptake (Maldonado et al., 2002; Rue and Bruland, 2001; Wells et al., 2005). DA production has also been shown to facilitate the competitive ability of *P. delicatissima* in co-culture with *S. marinoi* (Prince et al., 2013) supporting an allelopathic role (Xu et al., 2015). Alternatively, the change in allele frequency may be the result of other co-occurring biotic or abiotic drivers (including ocean circulation) that influence DA production but also exert a selective force on other species.

Whatever the reason, greater temporal and spatial resolution will help address these hypotheses. This can be accomplished by an increase in sequencing depth to obtain sufficient coverage for calling SNPs in an individual sample, which will increase per-sample sequencing costs. If the goal is to assess population level variation, it may be possible to enrich for the eukaryotic fraction by targeting not DNA, but RNA. Use of poly-A priming in the cDNA synthesis reaction can then be used to enrich for eukaryotes. This should not distort the relative abundance ratios of target taxa as the RNA to cDNA conversion does not involve any amplification, however, additional genotyping errors could be introduced during the conversion that could contribute to null alleles. Paired comparison of RNA and DNA based abundance estimates will be needed to evaluate the potential effectiveness of this modification.

### 4.2 Detection limit is driven by the false positive rate

The false positive rate greatly exceeded the false negative rate, but both were low on average, with less than 1% of reads identified as false positives in the mock community mix experiment. Although the method was able to detect the presence of species at very low abundances (tested down to 0.01%), an average false positive rate of 0.45% suggests that the detection limit for calling true positives should be set at 0.5-1%. Consequently, in the natural bloom sample set, although there were reads mapping to all species (Fig. 3q-t) only *P. australis, P. pungens*, and *P. subpacifica* were present at detectable levels. Interestingly, far fewer false positives were observed for reads mapping to the *P. multiseries* and *P. multistriata* reference species, which were *in silico* derived by extracting BcgI restriction sites from previously sequenced genomes. This despite *P. pungens* and *P. multiseries* being sibling species, which should increase overlap due to greater genetic similarity (Lim et al., 2018; Manhart et al., 1995). This suggests that some aspect of the culturing process may have contributed to artificially inflating the false positive rate. One explanation may be unintentional cross-contamination during the process of creating artificial community mixes, although the use of sterile technique makes this unlikely. Alternatively, false positives could still be the result of additional unknown contaminants (prokaryotic or eukaryotic), representatives of which have not yet been incorporated into the NCBI database. As 2b-RAD tags are only 36-bp in length, single base pair differences can be the difference between a match and a ‘no hits’ outcome. While we were able to identify and exclude the most contaminants using a global clustering approach, the vast majority of microbial diversity remains underexplored (Salazar and Sunagawa, 2017), and consequently cannot serve as a homology-based reference for filtering out contaminant reads. An alternative approach to excluding contaminants could be to focus not on homology based searches, which require a pre-existing reference database, but to use clustering algorithms, as have been developed for binning metagenomic sequence datasets using, for example tetranucleotide frequency and percent GC content (Graham et al., 2017). Future work should aim to explore if such alternative approaches can be applied to short read RAD datasets.

Accuracy should also be taken into consideration when setting detection limits. Cell size was not measured, but would be expected to vary considerably, along with genome size, across the different species utilized herein (Hubbard et al., 2014). It is possible that these differences contribute to variability in the expected vs. observed signal. In the mock community mix experiment, we observed significant variation in accuracy among replicates. Perhaps counter-intuitively, more equal mixes (e.g. ratios of 50:50) exhibited lower accuracy than more extreme mixes (e.g. ratios of 99.9:0.1) and pure cultures. In addition, accuracy varied significantly among replicate cultures of the same species. Taken together, this suggests that the variation in accuracy is likely driven by inaccuracies during mix creation rather than errors in the assignment of reads during mapping or systematic biases among species. As mixes were generated using replicate Sedgwick Rafter-based counts of cell densities in culture replicates, any inaccuracy during mix creation would propagate in proportion to the final volume of culture added to each sample mix. More equal mixes necessitated mixing of larger volumes for each individual culture, increasing the chance for variation. As it will be impossible in field samples to quantify the absolute abundance of any given focal species without a universal reference library, and given the strong association between observed and expected abundance (Fig. 2a), so long as the organism passes the minimum detection limit, then our results suggest its relative abundance within the focal community can be reliably assessed.

### 4.3. Low agreement of relative abundance estimates across methods

Both methods implicate *P. australis* as a driver of DA production in the natural sample set (Fig. 3q-t, Fig. 4) reinforcing earlier reports from this region *(Schnetzer et al*., *2013; Smith et al*., *2018b)*. However, direct comparison of the relative abundance estimates obtained using the different methods and even simple presence/absence determinations largely disagree. This discrepancy may be attributable to different methodological biases. After excluding samples below the 1% detection limit, 2bRAD still identified low abundances of *P. australis* (<5%) at almost all time-points (Fig. 3q-t) which were not evident in the ARISA analysis (Fig. 4). One explanation is false negatives in the ARISA analysis, as this method applies a 3% peak threshold. Lowering the ARISA limit of detection to 1% results in detection of *P. australis* in four additional samples, but a lowered threshold is not recommended without further quantitative validation. Alternatively, we cannot rule out the possibility that these low abundance detections are false positives in the 2bRAD analysis, as this method has a tendency to overestimate the abundance of species when rare, which is particularly evident in *P. australis* (Fig. 2a). Low agreement between methods is likely also driven by the limited diversity in the current 2bRAD reference library. The 2bRAD relative abundance estimates of all species will be artificially inflated by the inability to capture all the relevant diversity in *Pseudo-nitzschia* spp. Expansion of the reference library set to include additional relevant taxa for the region will help remedy this problem and shed more light on the ecology of this genus.

### 4.4 Proteobacteria: contaminants or symbionts?

Interactions between phytoplankton and bacteria are critical for nutrient cycling in aquatic environments, but recent work suggests that microscale interactions in the phycosphere, such as the exchange of metabolites, extend beyond general food web interactions into the realm of mutualism (Amin et al., 2012; Seymour et al., 2017). The essential relationship between diatoms and their associated bacteria may be why it is so difficult to obtain axenic phytoplankton cultures (Töpel et al., 2019). Bacteria have also been shown to enhance toxin production in *P. multiseries* cultures (Bates et al., 1995) suggesting a key role for other microbes in the ecology of *Pseudo-nitzschia* spp. We identified a substantial community of *Proteobacteria* in our unialgal *Pseudo-nitzschia* spp. cultures, the profiles of which differed among species. All cultures contained representatives of taxa previously identified as putative associates of diatoms and *Pseudo-nitzschia* spp. including *Sulfitobacter pseudonitzschiae* and *Marinobacter salarius* (Hong et al., 2015; Johansson et al., 2019) which occurred at low abundance (<5%) in most species. However, *P. australis* exhibited the highest level of ‘contamination’ with over 24% of the total RAD tags identified as *Proteobacteria* (Fig. S2) and of these contaminant tags, more than 93% exhibited blast matches to *M. salarius* in NCBI’s nt database. *M. salarius* has been shown to stimulate the growth of the diatom *Skeletonema marinoi* in culture (Johansson et al., 2019). Genomic analysis suggests that *M. salarius* may produce a growth factor in addition to siderophores, which may increase iron availability for its host diatom (Töpel et al., 2019). Future culture-based work should aim to test whether *M. salarius* fulfills a similar role for *P. australis*. From a methods development perspective, since 2bRAD captures both prokaryotic and eukaryotic tags, an interesting next step would be to expand the reference to include key prokaryotic taxa to assess the change in relative abundance of putative proteobacterial symbionts in addition to target *Pseudo-nitzschia* spp. over the course of natural bloom events.

### 4.5 Conclusions

Taken together, our results show that a reduced representation based sequencing approach can both quantify the relative abundance of *Pseudo-nitzschia* spp. in a mixed community and assess population-level genomic variation within species using the same dataset. Advantages of this method include simplicity of the library preparation protocol and cost-effectiveness (Puritz et al., 2014). Depending on the level of coverage desired, costs for both library preparation and sequencing range from ∼$10-$40 a sample, whereas full metagenomic libraries, which also require additional computational power to analyze, are on the order of $100 a sample. While other RAD methods could in principle be used, we advocate for 2bRAD for community level analyses as the fixed size of tags precludes the need for genome size corrections that would be necessary with other methods (Puritz et al., 2014), given that abundance is estimated from the total number of mapped reads, which is proportional to the per-taxa sequencing effort, rather than the mean coverage per tag. There are, however, some limitations that remain to be overcome. New species cannot be identified *a priori* in a mixed community. A RAD tag reference must first be generated. Culturing is a rate-limiting step for many methods, but with advances in single cell sequencing (Gawad et al., 2016), it may be possible to skip this step altogether in the near future in favor of cell sorting and direct sequencing (Cuvelier et al., 2010; Marie et al., 2017). By mining existing metagenomic datasets and other genome databases it may be possible to create a large-scale reference library that includes prokaryotes, archaea, and eukaryotes, to competitively recruit RAD tags from natural samples, thereby greatly expanding the taxonomic resolution of the method. Combined with the ability to secondarily assess population-scale variation for the most abundant members of the community, this method has the potential to greatly expand the scope of large-scale spatial and temporal monitoring studies, warranting additional development.

## Acknowledgements

We thank Carmelo Tomas for assistance in identification of *Pseudo-nitzschia* strains.

## Data Accessibility

FASTQ reads for both the Mock Community Mix and Natural Sample libraries can be obtained from NCBI’s SRA under BioProject PRJNA749297. The final *Pseudo-nitzschia* spp. 2bRAD reference library is hosted at https://dornsife.usc.edu/labs/carlslab/data/. Bioinformatic and statistical scripts necessary to re-create analyses, as well as raw input data used to generate figures used in this study are available at https://github.com/ckenkel/Pseudo-nitzschia2bRAD.

