## Supplementary Material for "Reduced representation sequencing accurately quantifies relative abundance and reveals population-level variation in *Pseudo-nitzschia* spp."

Figure S1. Scanning electron microscopy (SEM) images showing distinctive features of *Pseudo-nitzschia australis* (A) and *Pseudo-nitzschia pungens* (B) isolated from the Southern California Bight.

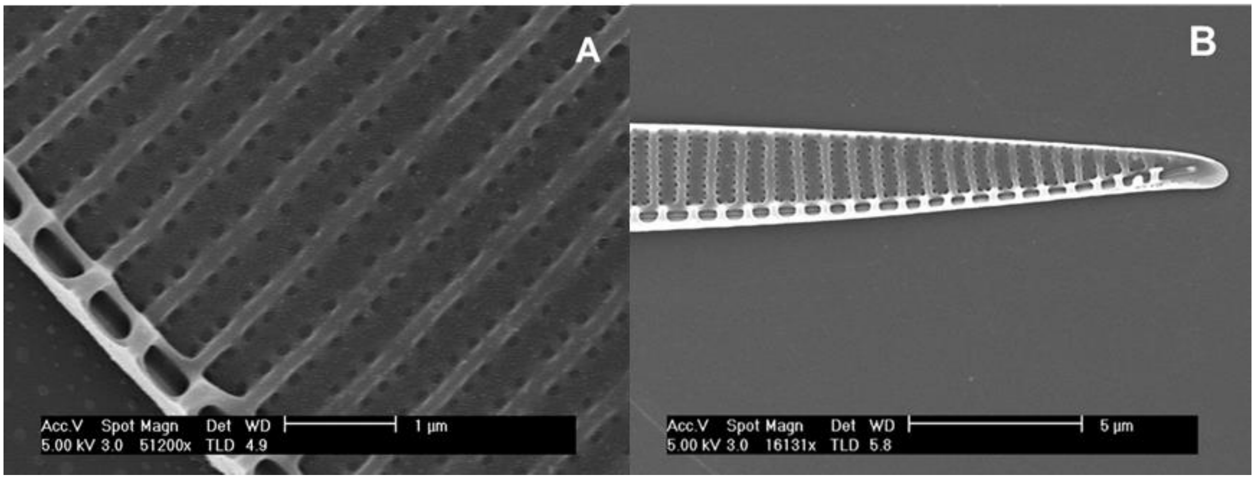

Figure S2. Proportion of tags within each reference species by taxonomic origin (as determined through a blastn search against the NCBI nt database).

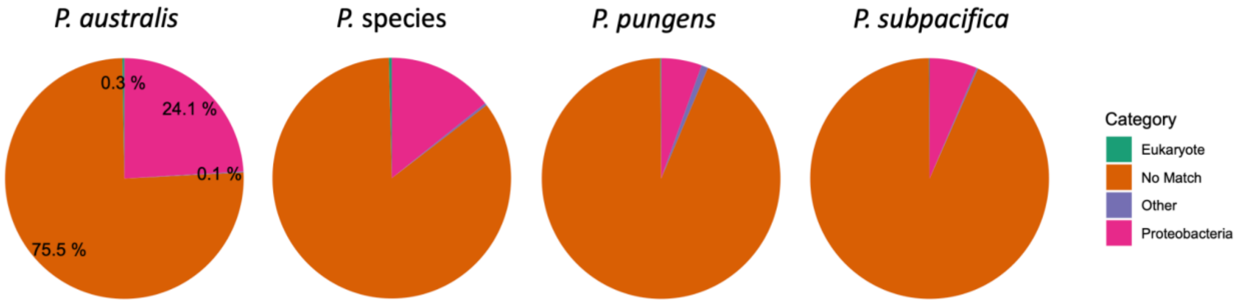


Figure S3. False positive rate by focal species and mix type.

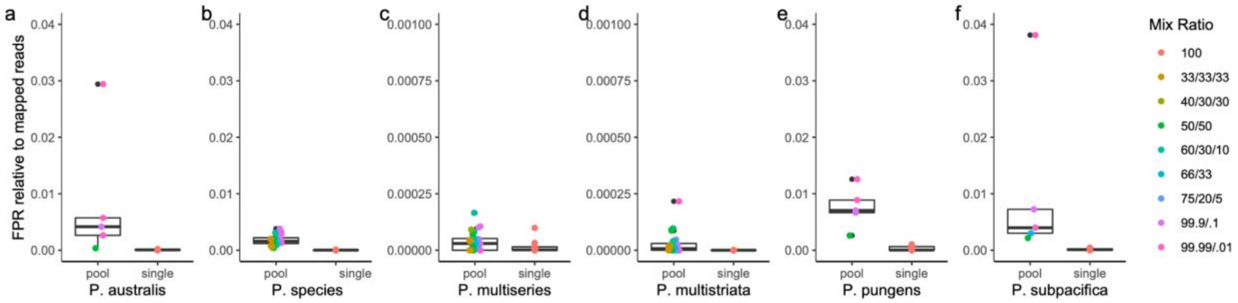


Figure S4. Domoic acid concentration (ng/mL) as a function of the relative percent abundance of reads mapping to *P. australis* across samples.

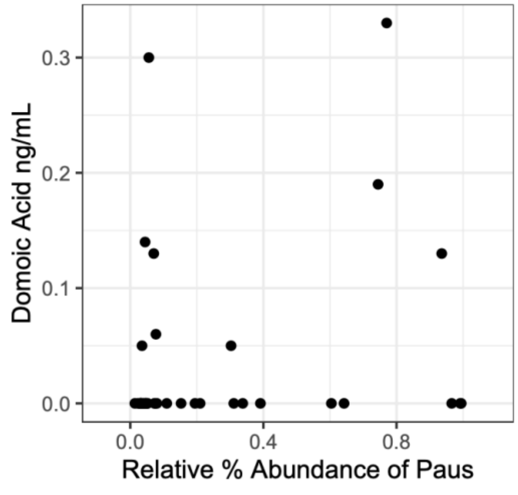

Figure S5. Percent of high quality reads exhibiting high quality mapping to any cluster in the reference library as a function of the relative percent abundance of reads mapping to *P. australis* across samples.

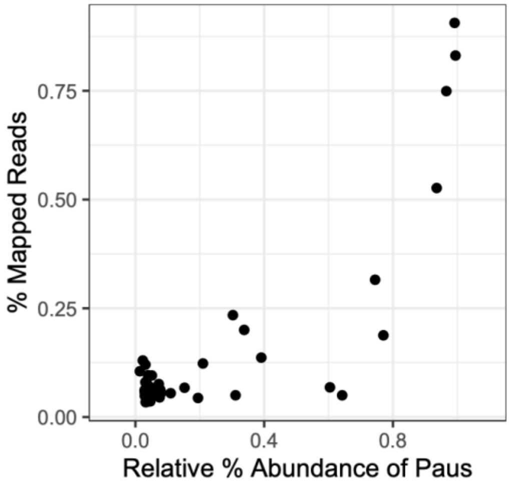


Figure S6. Relative abundance of *P. australis* estimated using the 2bRAD method as a function of the relative percent abundance of *P. australis*/*P. seriata* estimated using ARISA. The blue line indicates the actual correlation while the dashed line indicates 1:1.

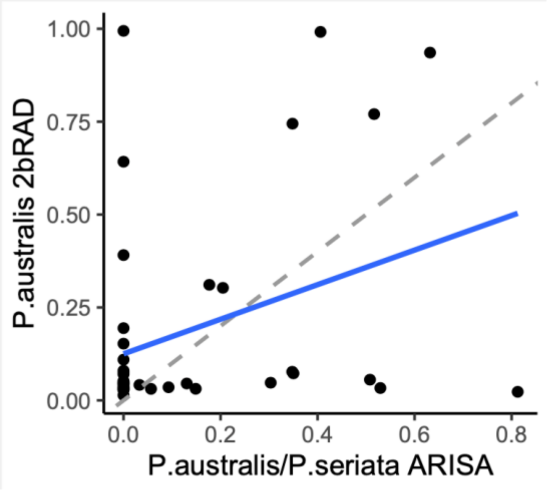

Figure S7. Density plot of the number of tags exceeding particular coverage thresholds for high quality reads exhibiting high quality mapping to the *P. pungens* reference.

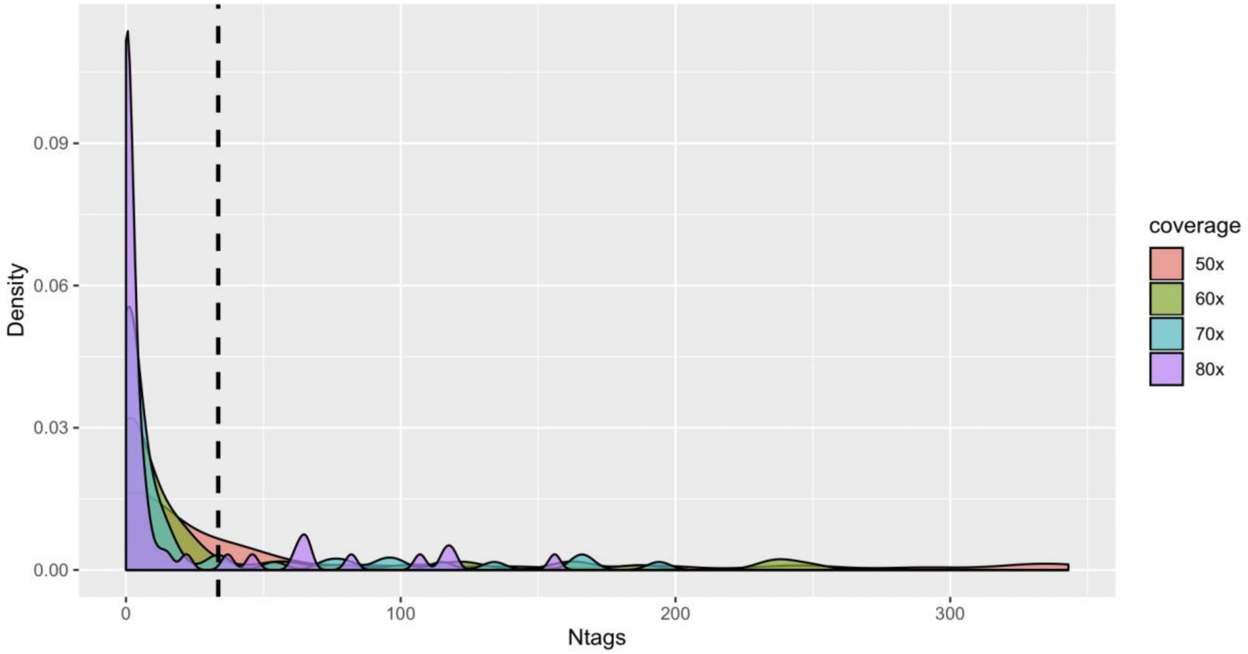


Table S1. Percent standard deviation for the triplicate counts of each of the *Pseudo-nitzschia* species used to create the mock community mixes shown in Table 1 of the main text.

| **Culture ID** | **Relative Standard Deviation** |
|---|---|
| *P. australis B6* | 10.0 |
| *P. pungens C3* | 2.0 |
| *P. pungens C5* | 5.9 |
| *P. pungens C6* | 4.5 |
| *P. subpacifica* | 7.1 |
